# Cadmium Disrupts Vestibular Function by Interfering with Otolith Formation

**DOI:** 10.1101/162347

**Authors:** Adrian J. Green, Carolyn J. Mattingly, Antonio Planchart

**Affiliations:** Department of Biological Sciences, North Carolina State University, Raleigh, NC; Center for Human Health and the Environment, North Carolina State University, Raleigh, NC

## Abstract

Cadmium (Cd^2+^) is a transition metal found ubiquitously in the earth’s crust and is extracted in the production of other metals such as copper, lead, and zinc^1,2^. Human exposure to Cd^2+^ occurs through food consumption, cigarette smoking, and the combustion of fossil fuels. Cd^2+^ has been shown to be nephrotoxic, neurotoxic, and osteotoxic, and is a known carcinogen. Animal studies and epidemiological studies have linked prenatal Cd^2+^ exposure to hyperactivity and balance disorders although the mechanisms remain unknown. In this study we show that zebrafish developmentally exposed to Cd^2+^ exhibit abnormal otolith development and show an increased tendency to swim in circles, observations that are consistent with an otolith-mediated vestibular defect, in addition to being hyperactive. We also demonstrate that the addition of calcium rescues otolith malformation and reduces circling behavior but has no ameliorating effect on hyperactivity, suggesting that hyperactivity and balance disorders in human populations exposed to Cd are manifestations of separate underlying molecular pathways.

## Introduction

The vestibular system of the vertebrate inner ear consists of three semicircular canals harboring sensory epithelia that respond to rotational acceleration. Two sensory organs, the utricle and saccule, which sense linear acceleration and gravity^3,4^ are located at the base of the semicircular canals. Together with the central nervous system, the vestibular system integrates both the magnitude and direction of head movements about an axis. The utricle and saccule house otoconia (termed otoliths in fish), which are bio-crystals composed of calcium carbonate (CaCO_3_) and proteins^3,4^, that lie above the ciliated hair cells of the sensory epithelium (macula); they are anchored to the hair cells by a honeycomb-like otoconial membrane^3^. If this membrane is compromised, a common occurrence in individuals with Ménière’s disease^5^, the otoconia can detach and dislocate causing vertigo and balance problems^3^. The correct formation and anchoring of otoconia are essential for optimal vestibular function, balance, and the detection of sound ^3,6,7^.

Zebrafish (*Danio rerio*) have become an increasingly valuable vertebrate model system in toxicology^41,15^ due to their high fecundity, external and transparent development, rapid generational times, reliance on evolutionarily conserved developmental regulatory gene networks, and sequenced genome, which facilitates mechanism-based toxicity studies. Like all vertebrate organisms, zebrafish have structural and sensory apparatus to perceive gravitational and positional cues for spatial awareness and posture control^16^, and perceive gravity and linear acceleration using mineralized biocrystals coupled to hair cells within the inner ear in a manner similar to other vertebrates, including humans.^16^ In zebrafish, these biocrystals are consolidated into three distinct and large structures termed otoliths that each directly interact with an entire patch of sensory epithelium located in the saccule, utricle, and a non-mammalian specific otolithic region known as the lagena^6^. The formation of otoliths in zebrafish has not yet been completely characterized but studies in trout and other species have shown that daily variations in the composition of vestibular endolymph are required ^8,9^. This diurnal variation in the composition of the endolymph consists of increasing the amount of protein and decreasing the concentration of calcium and bicarbonate ions in the endolymph during the day and reversing this process at night. This variation allows the otoliths to grow as the fish increases in size, which is a unique characteristic of fish not thought to occur in other vertebrates. However, many of the major proteins responsible for otoconial development in mammals are required for otolith formation in zebrafish, including the regulatory protein Otop1, which is expressed in hair cells during the seeding and growth of otoconia and otoliths and is responsible for regulating protein secretion and movement of calcium to and from the endolymph^3,18^. This anatomical and molecular conservation together with their transparency and experimental tractability make zebrafish a robust tool to identify mechanisms underlying environmentally induced ototoxicity.

Cadmium (Cd), is a non-essential transition metal widely used in the production of batteries, solar panels, pigments, plastic stabilizers, and the production of other metals ^10^. Oxidized cadmium (Cd^2+^) typically enters the body through contaminated food and water as well as through inhalation of polluted air and cigarette smoke^10,11^. Exposure to Cd^2+^ has been shown to cause nephrotoxic, neurotoxic, osteotoxic, and carcinogenic effects^10,11^, as well as ototoxicity (Agirdir et al. 2002; Kim et al., 2008, 2009; Ozcaglar et al., 2001; Liu et al. 2014). Epidemiological studies have linked prenatal Cd^2+^ exposure to hyperactivity and balance disorders^10^(Min et al., 2012), whereas i*n vivo* animal studies have shown that Cd^2+^ accumulates within the vestibular system and leads to defective hearing (Ozcaglar et al, 2001). Although the molecular mechanisms underpinning the vestibular defects observed in both epidemiological and animal studies remain unknown, one study demonstrated damage to hair cells resulting from exposure to Cd^2+^ (Liu et al., 2014). We hypothesized that exposure to Cd^2+^ alters the composition of the endolymph leading to the production of abnormal otoconia/otoliths, which results in disruption of the otoconia/otolith-hair cell interaction, thus causing disorders characteristic of vestibular defects. In this study we show that zebrafish exposed developmentally to Cd^2+^ exhibit hyperactivity, circling behavior, altered otolith development and hair cell interactions, and have disrupted *otop1* gene expression. Furthermore, supplementation with calcium rescues otolith formation and diminishes associated circling behavior but has no effect on the observed hyperactivity. These findings suggest that behavioral abnormalities resulting from Cd toxicity can be subdivided into disorders rooted in ototoxicity and disorders rooted in neurological impairment, each having a distinct mechanistic explanation.

## Materials and Methods

### Animal Husbandry

Wildtype (AB strain) zebrafish were maintained at the NC State University Zebrafish Core Facility according to standard protocols, including 14-10 light cycle, *ad libitum* feeding three times daily, and maintenance at 28.5° C ^20^. All work involving zebrafish was approved by the NC State Animal Care and Use Committee.

### Chemicals

Stock solutions of cadmium chloride (Cd^2+^, Cat. # Sigma-Aldrich, MO), N-Acetyl Cysteine (NAC, Cat. # Sigma-Aldrich, MO), and calcium chloride (Cat. # Sigma-Aldrich, MO) were made in reagent-grade (Picopure^®^) water at 10 parts-per-thousand (CdCl_2_), 25 mg/mL (NAC) and 0.5 M (CaCl_2_), and stored at room temperature in 1.5 mL polypropylene tubes. 10X embryo media (E2), consisting of 150 μM NaCl, 5 μM KCl, 10 μM MgSO_4_, 1.5 μM KH_2_PO_4_, 0.5 μM Na_2_HPO_4_, 10 μM CaCl_2_, and 7 μM NaHCO_3_ was prepared in reagent-grade water and diluted to 0.5X for subsequent exposure studies.

### Exposures

Zebrafish embryos were collected immediately after spawning and exposed to 30 - 60 parts-per-billion (ppb) Cd^2+^ in 0.5X E2 media from four hours post-fertilization (hpf) through seven days post-fertilization (dpf) at a density of 10 embryos/mL. This concentration range was selected as it represents the upper range of exposure observed in human populations exposed to cadmium-polluted environments^2,11,21,22^. The media was replaced daily and feeding began at five dpf. Cadmium has been shown to induce oxidative stress and interfere with ion channels, in particular zinc and calcium channels^24,25^. To evaluate whether these effects were the primary cause of the observed Cd-induced otolith and/or behavioral phenotypes, embryos were co-exposed from four hpf to 7 dpf with daily media changes to Cd^2+^ (60 ppb) and either 1 mg/L N-acetyl-L-cysteine (NAC) or 2.5 mM Calcium (CaCl_2_).

### Otolith Imaging and Measurements

Light microscopy images of otoliths were taken using a Leica dissecting microscope. To assess otolith size, seven dpf zebrafish larvae (n=5) were anesthetized by the addition of phosphate-buffered Tricaine (final concentration: 0.4 mg/L, pH 7, Western Chemical Inc.) to 0.5X E2 media until no movement was observed. Once anesthetized, images were taken using the Leica Application Suite (software version 4.8.0). The length of the otoliths, which resemble prolate spheroids, was determined by measuring across the longest axis using the GNU Image Manipulation Programs protractor tool (version 2.8.18)^23^.

### Scanning Electron Microscopy

Seven dpf zebrafish larvae were pooled in 1.5 mL polypropylene tubes in triplicate (n=10 per replicate) and placed on ice for 10-15 minutes to euthanize them. Larvae were dehydrated using successive 15-minute incubations in a graded ethanol series (33%, 66%, and 100%), and otoliths were dissected using fine-tipped forceps under a Leica dissecting microscope and stored in 100% ethanol at 4°C for a maximum of seven days. Otoliths were fixed in 3% glutaraldehyde in 0.1M NaPO_4_ buffer (phosphate buffer), pH 7.4 at 4°C and afterwards transferred to microporous specimen capsules (Structure Probe Inc., West Chester PA) for processing. Samples were washed in three changes of phosphate buffer and dehydrated in a graded ethanol series to 100% before critical point drying in liquid CO_2_ (Tousimis Research Corporation, Rockville MD). Otoliths were placed on carbon tabs (Ladd Research Industries, Williston VT) and sputter coated with Au/Pd using a Hummer 6.2 sputter system (Anatech USA, Union City CA). Samples were viewed on a JEOL JSM-5900LV at 15kV (JEOL USA, Peabody MA).

### Behavioral Assays

Behavior was assessed using a DanioVision™ box with EthoVision^®^ XT software (Noldus; Leesburg, VA). Behavior analyses for all exposures were conducted at five dpf. Zebrafish larvae were arrayed in a six-well plate at a density of eight larvae per well. Plates were placed in the DanioVision™ box and larvae were allowed to acclimate in the dark for at least 30 minutes. Following acclimation, larval response to three 10- minute dark-light cycles was measured. Measured end points included movement, acceleration, velocity, and rotation, including binning into clockwise or counterclockwise rotation. As zebrafish larvae are more active in the dark, cumulative distance moved in the dark was used to compare treated vs. controls.

### Quantitative PCR

Larvae (n=10 per replicate) were pooled in triplicate at 6, 12, 18, 24, 36, 48, and 120 hpf into 1.5 mL polypropylene tubes and euthanized by placement on ice for 10 minutes. All liquid was removed and 800 μL of TriReagent (Sigma-Aldrich, MO) was added to each tube. Larvae were homogenized using RNase-free disposable micropestles. Total RNA was extracted according to the manufacturer’s protocol and the integrity and concentration were assessed on an Agilent Bioanalyzer using the Agilent RNA 6000 Nano Kit (Agilent Technologies, CA). Equal amounts of total RNA from each sample were used to make cDNA by reverse transcription PCR using a poly-dT primer. Quantitative PCR was performed using SYBR Green and ROX reference dye (Agilent Technologies, CA). Primer sequences for star marker (*stm*), secreted protein, acidic, cysteine-rich (*sparc*), otopetrin 1 (*otop1*), and eukaryotic translation elongation factor 1 alpha 1, like 1 (*eef1a1l1*) are provide in Table 1. All qPCR assays were normalized to *eef1a1l1*, which did not vary across treatment groups. Primer pairs were designed to exons separated by at least one intron to avoid amplifying genomic DNA. The 2^(- ΔΔCT) method was used to approximate relative transcript fold change between treatment groups ^24^.

**Table 1.**
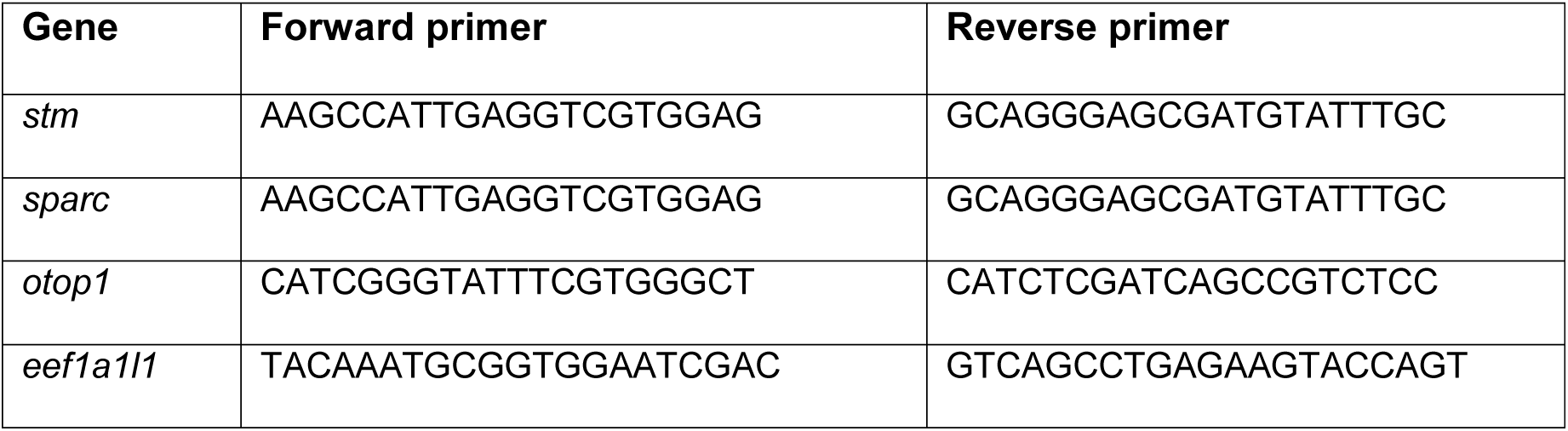
PCR Primer Sequences Used for Qpcr.

## Results

### Cadmium Induces Changes in Otolith Diameter

Developmental cadmium exposure resulted in a decrease in the diameter of the saccule otolith at 7 dpf relative to untreated controls (Figure 1); however, the utricle otolith diameter was not significantly different between controls and Cd-exposed larvae. The decrease in saccule otolith also exhibited a dose-dependent decrease beginning at 30 ppb (p < 0.01) and leveling off at 40 ppb cadmium (p < 0.001) (Fig. 1d). These results demonstrate that developmental Cd^2+^ exposure reduces the size of the saccule otolith in larval zebrafish without affecting the utricle otolith.

**Figure 1:**
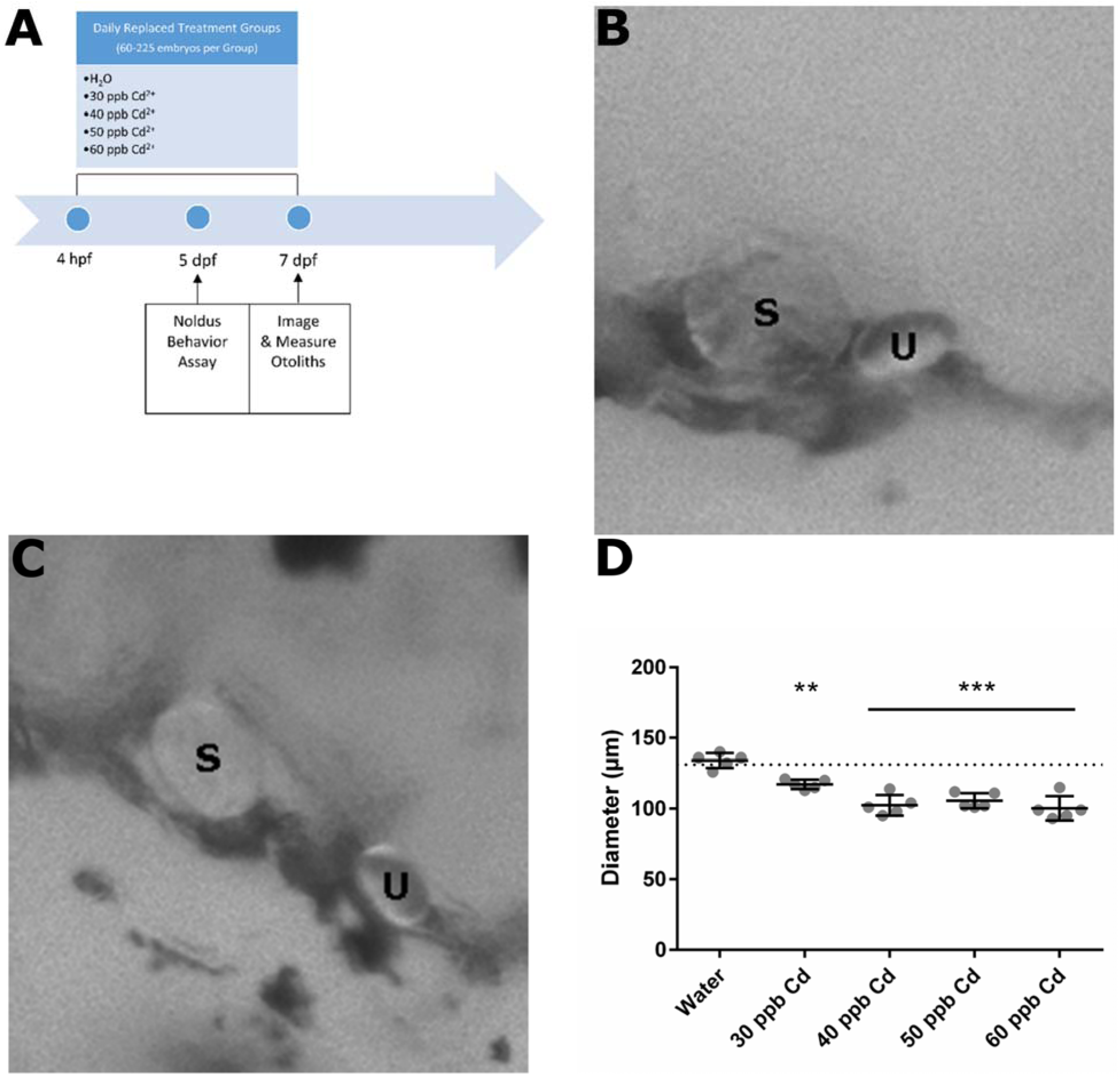
Otolith Size Decreases in Response to Cd^2+^ Exposure. (A) Zebrafish dose response to an exposure of 30 – 60 parts per billion (ppb) Cadmium (Cd^2+^). (B) Unexposed zebrafish otolith at seven dpf. (C) Zebrafish otolith exposed to 60 ppb Cd^2+^ at seven dpf. (D) Diameter of zebrafish saccule otolith at seven dpf. saccule otolith (S), utricle otolith (U); **p < 0.01, ***p< 0.001.

### Cadmium Alters Otolith Ultrastructure

To assess the ultrastructure of the saccule otoliths, we used scanning EM (SEM) to visualize control and Cd-exposed saccule otoliths. Our initial analysis revealed pronounced structural differences in the Cd-exposed otoliths compared to control otoliths. Whereas, control otoliths exhibited a smooth surface with evidence of otolith attachment to hair cell stereocilia, Cd-exposed otoliths exhibited a course knobbled surface with numerous fiber or cable-like extensions between knobs and an absence of evidence of attachment to hair cells (Fig. 2). The presence of numerous cable-like extensions and the pronounced increase in surface area due to surface knobbling in the Cd-exposed otoliths suggests a defect in the deposition of inorganic minerals or specific proteinaceous components required for normal otolith formation.

**Figure 2:**
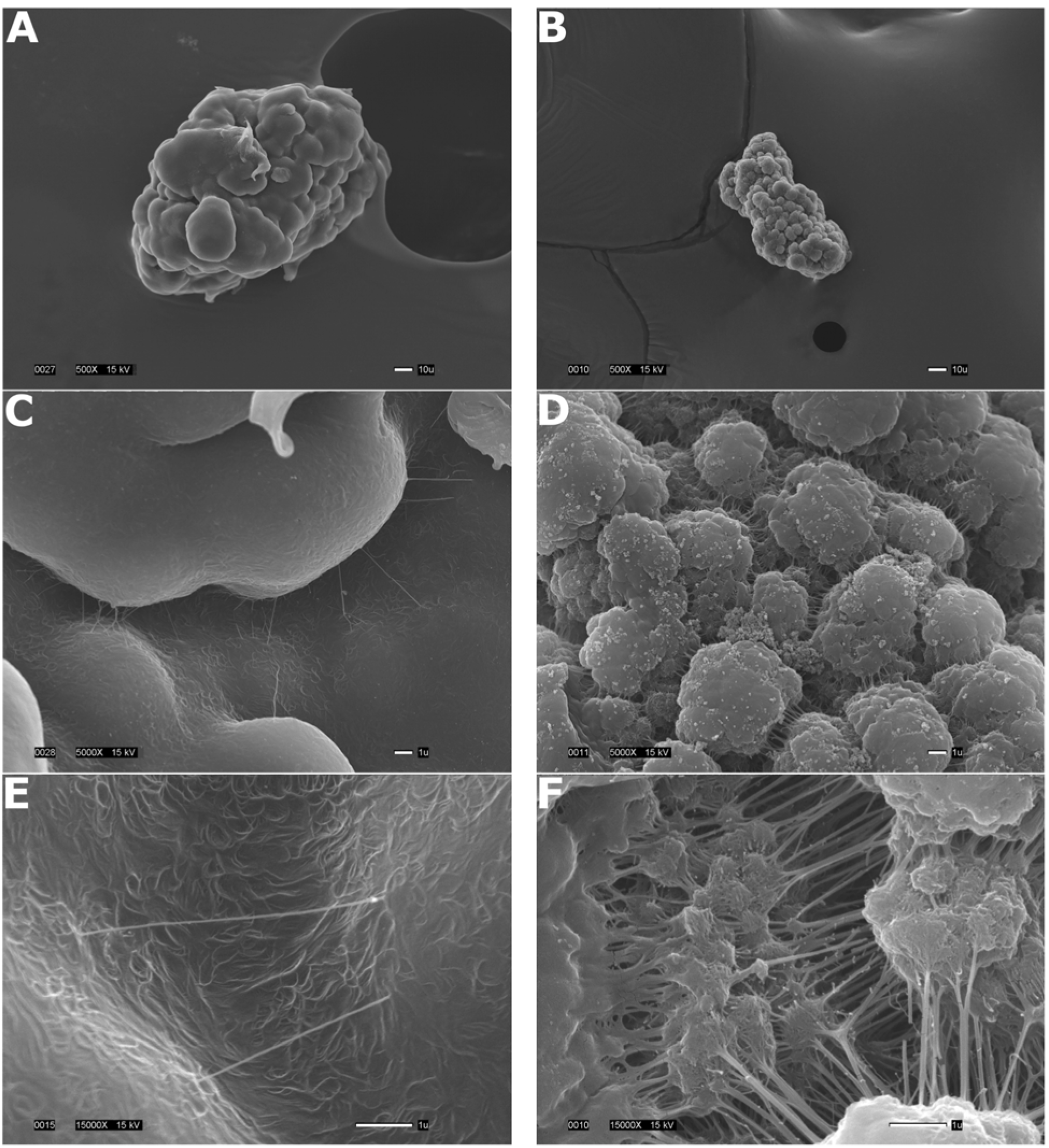
Saccule Otolith SEM Images at Seven dpf. A) Unexposed zebrafish otolith. (B) Zebrafish otolith exposed to 60 ppb Cd2+. (C) High magnification image of unexposed otolith. (D) High magnification image of otolith exposed to 60 ppb Cd2+. (E) Very high magnification image of unexposed otolith. (F) High magnification image of otolith exposed to 60 ppb Cd2+. Study design for the collection for zebrafish saccule otolith to assess Cd2+ induced changes in ultrastructure using scanning electron microscopy (SEM).

### Cadmium Induces Behavioral Changes Associated with Vestibular Defects

To determine if the otolith phenotype observed in Cd-exposed zebrafish correlated with changes in behavior, we assessed several behavioral endpoints, including activity in light versus dark, and tendency to swim in circles. Cadmium caused a dose-dependent increase in distance moved during dark periods consistent with a hyperactive-like phenotype. As seen in Figure 3, zebrafish larvae exposed to 40 - 60 ppb cadmium exhibited a heightened level of activity throughout the behavioral assay but this difference was most pronounced during dark periods (p < 0.05) when zebrafish larvae are typically more active^25,26^. In addition to increased activity in the dark, Cd^2+^ increased the number of rotations in both a clockwise (CW) and counter clockwise (CCW) direction but with a clear preference for CCW movement (p < 0.0001; See Supplemental Video). This circling behavior is characterized by a rapid twirling or spinning behavior in response to both mechanical and light stimuli and is often observed in other model organisms harboring mutations that affect the vestibular system^27,28^. These data demonstrate that Cd^2+^ induces a dose-dependent increase in activity and number of rotations in 5 dpf larval zebrafish.

**Figure 3:**
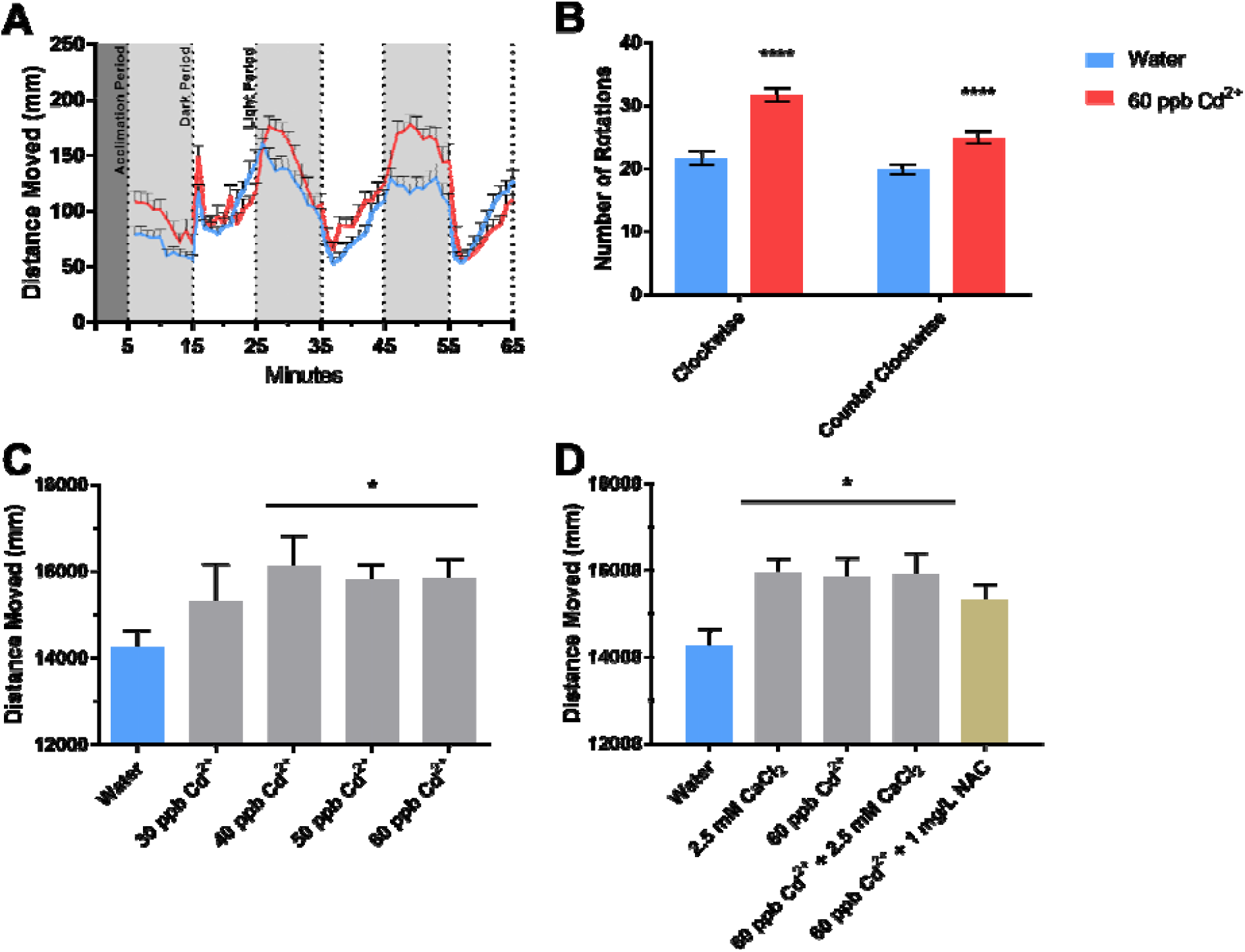
Cd^2+^ Induced Hyperactivity and Rotations. (A) Zebrafish distance moved in response to light and dark cycling. (B) Number of clockwise and counter clockwise rotations in the dark in response to Cd. (C) Average total distance moved in the dark in response to Cd. (D) Average total distance moved in the dark in response to rescue treatments. (A-D) Behavioral assessments made using Noldus Ethovision software; * p < 0.05, **** p < 0.0001.

### Calcium Rescues Otolith Formation and Diminishes Vestibular-Related Rotational Behavior but not Hyperactivity

Cadmium has been shown to interfere with divalent ion channel function, in particular zinc channels and both voltage and non-voltage gated calcium channels, and to induce oxidative stress^29,30^. To test the hypothesis that Cd^2+^ interferes with calcium homeostasis in the vestibular system, zebrafish embryos were co-exposed to Cd^2+^ and the effect of exogenous calcium on otolith formation was assessed. As seen in Figure 4, calcium prevented Cd-mediated decrease in saccule otolith size. In addition, the inclusion of calcium in the media was associated with a decrease in the rotational behavior observed in presence of Cd^2+^ only, indicating that the defects in otolith formation were inducing behavioral effects commonly observed in other model organisms with vestibular defects. However, the addition of calcium was not effective at reducing hyperactivity (Fig 4). In fact, addition of calcium alone induced a hyperactive phenotype independent of Cd^2+^ exposure. To test the hypothesis that changes in otolith size and hyperactivity were due to Cd^2+^-dependent increases in reactive oxygen species (ROS), a phenomenon reported in numerous studies^13^, we exposed larvae to Cd^2+^ alone or in combination with NAC, a ROS scavenger, Whereas NAC was able to rescue hyperactivity, it did not prevent the decrease in saccule otolith size nor the increased circling behavior induced by Cd exposure (Fig 4). These results provide evidence that Cd-induced hyperactivity and circling behaviors are independent events likely due to the disruption of different molecular pathways, including vestibular (circling) and neurological (hyperactivity) pathways, and that hyperactivity is not dependent on otolith malformations and associated vestibular defects.

**Figure 4:**
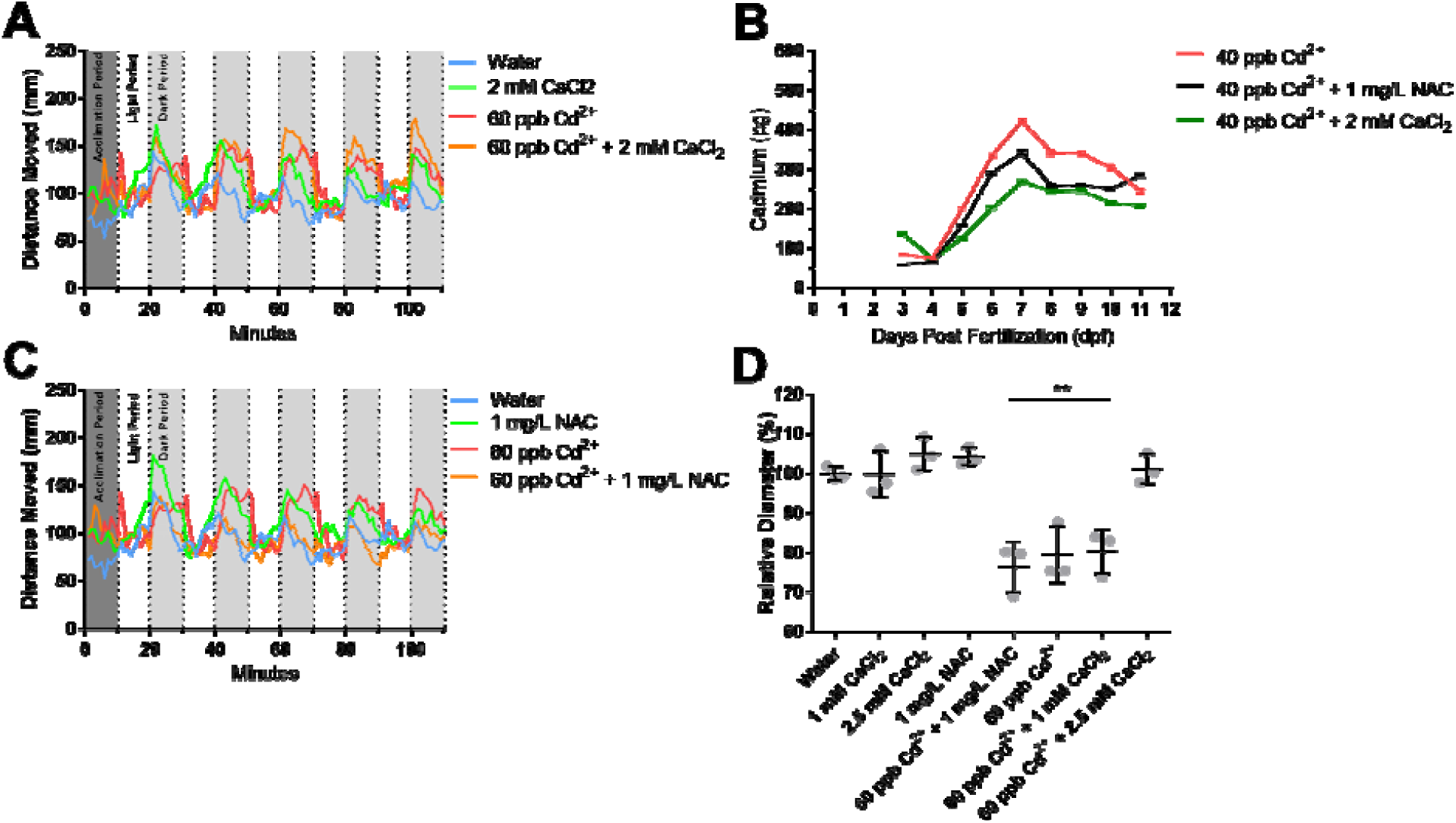
Otolith Recovers when Treated with Ca. (A) Otolith rescue experiments using 1 mg/L N-Acetyl Cysteine (NAC) or 2.5 mM Calcium. (B) Changes in saccule otolith diameter relative to water controls with rescue treatments; ** p < 0.01.

### Cadmium Induces Changes in mRNA expression of the Otolith Protein, *otop1*

Analysis of genes important for otolith development revealed that expression of the calcium regulatory gene *otop1* is significantly affected by exposure to Cd^2+^ during early development. At 6 and 24 hpf, *otop1* gene expression was significantly (p < 0.05) reduced in Cd-exposed versus unexposed embryos (Fig 5). This trend reversed at 36 hpf at which point it was significantly (p < 0.05) elevated in Cd-exposed embryos. Otolith genes that encode for the structural proteins Sparc, and Stm were not significantly altered at any time points tested. These data demonstrate that zebrafish developmentally exposed to Cd^2+^ exhibit mRNA expression changes in *otop1*, a gene important for vestibular calcium sensing and signaling.

**Figure 5:**
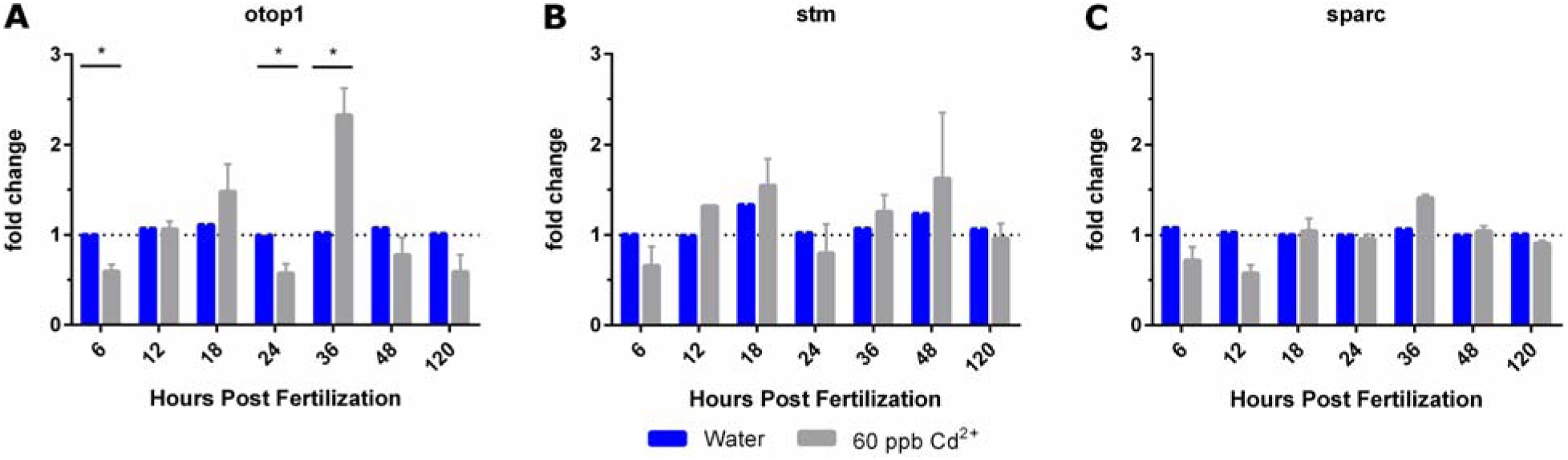
Quantitative PCR of Otolith Genes from 4 - 120 Hours Post-Fertilization. (A) Otopetrin 1 (otop1) gene expression. (B) Starmaker (stm) gene expression. (C) Secreted protein, acidic, cysteine-rich (sparc) gene expression; *p < 0.05.

## Discussion

Developmental exposure to Cd^2+^ reduces the size of the saccule otolith in a dose-dependent manner. This decrease was characterized by a 21% reduction in the size of the saccule otolith (Fig 1) at 40, 50, and 60 ppb Cd^2+^. SEM revealed that not only was the saccule otolith reduced in size but that there are significant changes in the ultrastructure. Otoliths exposed to Cd^2+^ have a rough cauliflower surface texture whereas the control otoliths are relatively smooth (Fig 2). We observe that Cd^2+^induced hyperactivity in a dose-dependent manner by assessing average total distance moved in the dark (Fig 3) and that calcium is capable of rescuing otolith formation but not hyperactivity (Fig 3, Fig 4). Calcium alone is capable of inducing hyperactivity which might account for why it only rescues otolith formation. NAC was able to reduce cadmium-induced hyperactivity but could not rescue otolith formation (Fig 3, Fig 4)

Semi-quantitative qPCR analysis revealed that Cd^2+^ altered the expression level of *otop1*, a gene important for vestibular calcium sensing and transport. More work is needed to determine whether or not the Cd^2+^-induced reduction in saccule otolith size is responsible for the observed hyperactivity and/or rotation phenotype. The ultrastructure showed that the Cd^2+^-exposed saccule otoliths were missing structural elements and when combined the observations of changes in *otop1* gene expression hints at a possible loss of calcium deposits. Further the cable-like structure seen in the SEM images may be a type of structural or scaffolding element required during the biosynthesis of the otoliths. To help answer these questions we intend to test additional ROS scavengers^31^, evaluate the uptake of Cd^2+^ using a Cd-109 radiotracer in the presence and absence of potentially protective compounds such as ebselen, calcium, and ascorbic acid, determine the protein and calcium composition of the saccule otoliths using Inductively coupled plasma mass spectrometry (ICP-MS), and determine the mechanism by which Cd^2+^ disruptions otop1 and its effects on otolith development.

Hyperactive behavioral disorders were estimated to affect between 5-11% for children between 4-17 years of age in 2011 (6.4 million)^32^. The data we have presented provides a plausible mechanistic link between an inner ear defect and hyperactivity because of Cd^2+^-exposure. By using common fairly innocuous compounds, as demonstrated here, it may be possible to reduce the number of children born with ADHD in the future.

